# mRNA bivalent booster enhances neutralization against BA.2.75.2 and BQ.1.1

**DOI:** 10.1101/2022.10.31.514636

**Authors:** Meredith E. Davis-Gardner, Lilin Lai, Bushra Wali, Hady Samaha, Daniel Solis, Matthew Lee, Andrea Porter-Morrison, Ian Thomas Hentenaar, Fumiko Yamamoto, Sucheta Godbole, Daniel C. Douek, Frances Eun-Hyung Lee, Nadine Rouphael, Alberto Moreno, Benjamin A. Pinsky, Mehul S. Suthar

**Affiliations:** Emory University School of Medicine; Atlanta, GA; Stanford University School of Medicine; Stanford, CA; National Institute of Allergy and Infectious Diseases; Bethesda, MD

**Author notes:** Correspondence: Mehul S. Suthar.

## Abstract

The emergence of the highly divergent SARS-CoV-2 Omicron variant has jeopardized the efficacy of vaccines based on the ancestral spike. The bivalent COVID-19 mRNA booster vaccine within the United States is comprised of the ancestral and the Omicron BA.5 spike. Since its approval and distribution, additional Omicron subvariants have been identified with key mutations within the spike protein receptor binding domain that are predicted to escape vaccine sera. Of particular concern is the R346T mutation which has arisen in multiple subvariants, including BA.2.75.2 and BQ.1.1. Using a live virus neutralization assay, we evaluated serum samples from individuals who had received either one or two monovalent boosters or the bivalent booster to determine neutralizing activity against wild-type (WA1/2020) virus and Omicron subvariants BA.1, BA.5, BA.2.75.2, and BQ.1.1. In the one monovalent booster cohort, relative to WA1/2020, we observed a reduction in neutralization titers of 9-15-fold against BA.1 and BA.5 and 28-39-fold against BA.2.75.2 and BQ.1.1. In the BA.5-containing bivalent booster cohort, the neutralizing activity improved against all the Omicron subvariants. Relative to WA1/2020, we observed a reduction in neutralization titers of 3.7- and 4-fold against BA.1 and BA.5, respectively, and 11.5- and 21-fold against BA.2.75.2 and BQ.1.1, respectively. These data suggest that the bivalent mRNA booster vaccine broadens humoral immunity against the Omicron subvariants.

## Results

The emergence of the highly divergent Omicron variant of SARS-CoV-2 led to concerns about the efficacy of vaccines based on the ancestral spike, and the approval of bivalent COVID-19 vaccines within the United States (the ancestral spike and the Omicron subvariant BA.5 spike proteins)^1–4^. Since its approval and distribution, additional subvariants have been identified with key mutations that further escape vaccine-elicited antibodies and approved monoclonal antibodies^5^. Of particular concern is the R346T mutation which has arisen in multiple variants of different lineages, including BA.2.75.2 and BQ.1.1 (**Supplementary Fig. 1**). We tested serum samples from individuals who had received either one or two monovalent boosters or the bivalent booster to determine neutralization efficiency against wild-type (WA1/2020) virus and Omicron subvariants BA.1, BA.5, BA.2.75.2, and BQ.1.1 using a live virus neutralization assay.

We used an *in vitro*, live-virus focus neutralization test (FRNT) assay in a VeroE6-TMPRSS2 cell line^1^ to compare the neutralizing activity of serum from individuals who received one monovalent booster (7-28 days after vaccination), two monovalent boosters (70-100 days after vaccination), or the bivalent booster (16-42 days after vaccination). The fold change in neutralizing antibody response among these three cohorts were quantitated by comparing the FRNT_50_ GMT (geometric mean titer) values of Omicron against the ancestral SARS-CoV-2 virus. Samples that fell below the limit of detection (1:20) were given an arbitrary FRNT_50_ of 10.

In all groups, a decrease in neutralization activity was observed against all omicron subvariants compared to WA1/2020, with the greatest decrease seen against BQ.1.1 (**Fig. 1**). In the one monovalent booster cohort, the FRNT_50_ GMTs were 758 for WA1/2020, 60 for BA.1, 50 for BA.5, 23 for BA.2.75.2 and 19 for BQ.1.1. In the two monovalent booster cohort, the FRNT_50_ GMTs were 1812 for WA1/2020, 205 for BA.1, 142 for BA.5, 65 for BA.2.75.2 and 53 for BQ.1.1. In both cohorts, relative to WA1/2020, this corresponded to a reduction in neutralization titers of 9-15 fold against BA.1 and BA.5 and 28-39 fold against BA.2.75.2 and BQ.1.1. BA.2.75 showed comparable neutralization titers as BA.1 and BA.5 in these cohorts (**Supplemental Fig. 2**).

**Figure 1.**
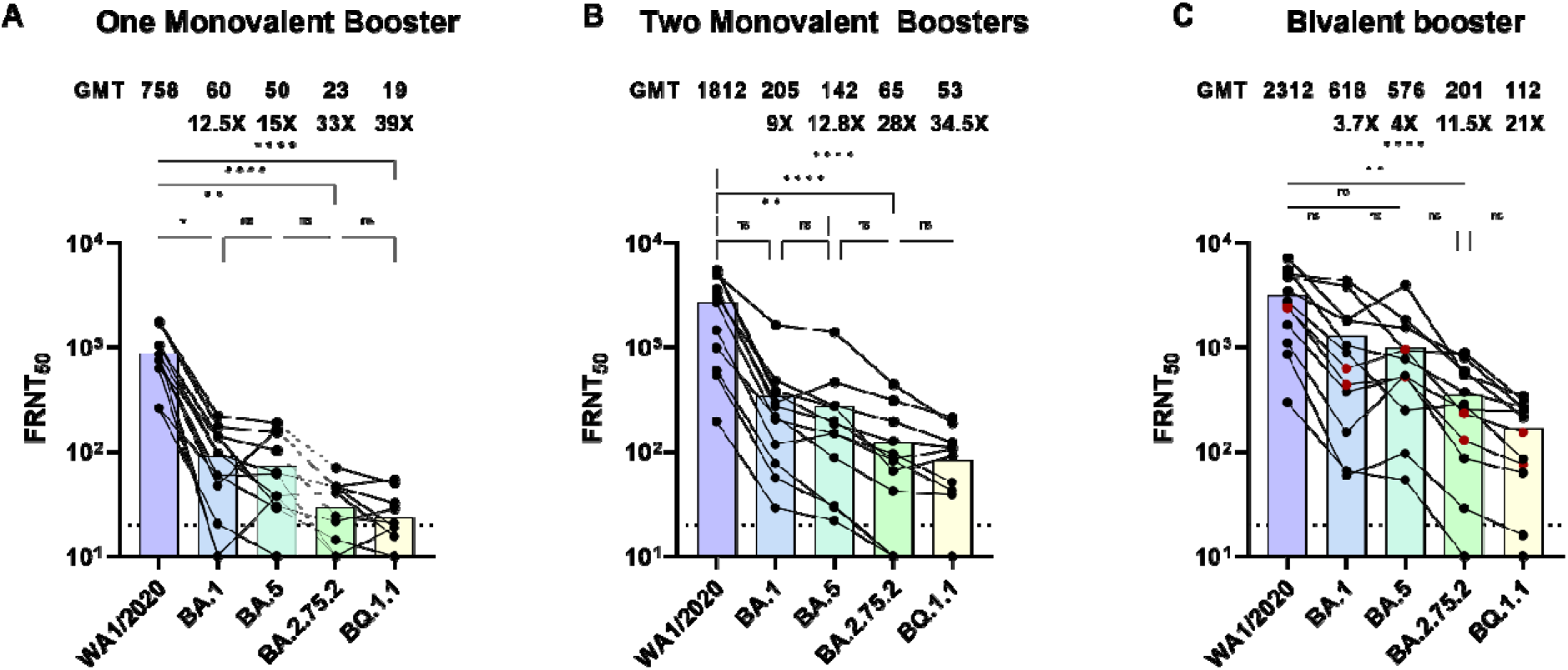
Neutralizing responses against WA1/2020, BA.1, BA.5, BA.2.75.2, and BQ.1.1. Shown is the neutralization activity against SARS-CoV-2 variants among 12 individuals who received one monovalent booster (Panel A), 12 individuals of received two monovalent boosters (Panel B), and 12 individuals who received the updated bivalent booster (Panel C). The focus reduction neutralization test (FRNT_50_ [the reciprocal dilution of serum that neutralizes 50% of the input virus]) geometric mean titers for each variant are shown above each panel along with ratios of GMT compared to WA1/2020. The connecting lines between the variants represent matched serum samples. The horizontal lines represent the limit of detection of the assay (FRNT_50_ GMT 20). Red symbols in panel C indicate two individuals self-reported prior SARS-CoV-2 infection. The differences between all groups were determined with the Kruskal–Wallis test with Dunn’s correction for multiple comparisons. * p<0.05, ** p<0.01, ***p<0.001, **** p<0.0001, n.s not significant.

In the BA.5-containing bivalent booster cohort, the neutralizing activity improved against all of the Omicron subvariants (**Fig 1C**). The FRNT_50_ GMTs were 2312 for WA1/2020, 618 for BA.1, 576 for BA.5, 201 for BA.2.75.2 and 112 for BQ.1.1. Relative to WA1/2020, this corresponded to a reduction in neutralization titers of 4-fold against BA.1 and BA.5 and 11- and 21- fold against BA.2.75.2 and BQ.1.1, respectively.

Individuals that received either one or two monovalent COVID-19 boosters had a dramatic decrease in neutralization activity against Omicron subvariants compared to WA1/2020. This decrease was especially profound for BA.2.75.2 and BQ.1.1, which contain the predicted escape mutation R346T. Individuals that received the BA.5-containing bivalent booster showed improved neutralizing activity against all Omicron subvariants. These responses are similar to recent observations in individuals with breakthrough Omicron infection showing broadened neutralizing activity against Omicron variants^6^. Limitations of this study include small cohort size, unknown impact of prior SARS-CoV-2 exposure, and examination of a single timepoint. These data demonstrate an overall serological benefit of bivalent booster immunizations.

