## Supplementary Fig. 1 for "mRNA bivalent booster enhances neutralization against BA.2.75.2 and BQ.1.1"

**Supplementary Appendix**

### Supplemental Methods

**Serum samples**. For samples Emory University, collection and processing were performed under approval from the University Institutional Review Board (#00002061 and #00058271). Adults ≥18 years were enrolled who met eligibility criteria under these protocols, were at least 1 week post-COVID booster vaccination, and provided informed consent. All patient samples were de-identified prior to inclusion in the study.

**Cells and Viruses.** Vero-TMPRSS2 cells were cultured in complete DMEM medium consisting of 1x DMEM (VWR, #45000-304), 10% FBS, 2mM L-glutamine, and 1x antibiotic as previously described^1^. nCoV/USA_WA1/2020 (WA/1), closely resembling the original Wuhan strain, was propagated from an infectious SARS-CoV-2 clone as previously described^2^. icSARS-CoV-2 was passed once to generate a working stock. The BA.1 isolate has been previously described (Edara Cell Reports 2022). Omicron subvariants were isolated from residual nasal swabs: BA.5 isolate (EPI_ISL_13512579), provided by Dr. Richard Webby (St Jude Children’s Research Hospital), BA.2.75.2 (EPI_ISL_15146622), BQ.1.1 isolate (EPI_ISL_15196219), and BA.2.75 isolates (EPI_ISL_14393635) provided by Dr. Benjamin Pinsky (Stanford University). All variants were plaque purified and propagated once in VeroE6-TMPRSS2 cells to generate working stocks.

**Focus Reduction Neutralization Assay.** FRNT assays were performed as previously described^3^. Briefly, samples were diluted at 3-fold in 8 serial dilutions using DMEM in duplicates with an initial dilution of 1:10 in a total volume of 60 μl. Serially diluted samples were incubated with an equal volume of SARS-CoV-2 (100-200 foci per well) at 37^o^ C for 1 hour in a round-bottomed 96-well culture plate. The antibody-virus mixture was then added to Vero cells and incubated at 37^o^ C for 1 hour. Post-incubation, the antibody-virus mixture was removed and 100 µl of prewarmed 0.85% methylcellulose overlay was added to each well. Plates were incubated at 37^o^ C for 18 to 40 hours, and the methylcellulose overlay was removed and washed six times with PBS. Cells were fixed with 2% paraformaldehyde in PBS for 30 minutes. Following fixation, plates were washed twice with PBS, and permeabilization buffer (0.1% BSA, 0.1% Saponin in PBS) was added to permeabilize cells for at least 20 minutes. Cells were incubated with an anti-SARS-CoV spike primary antibody directly conjugated to Alexa Fluor-647 (CR3022-AF647) overnight at 4°C. Cells were then washed twice with 1x PBS and imaged on an ELISPOT reader (CTL Analyzer).

**Quantification and Statistical Analysis.** Antibody neutralization was quantified by counting the number of foci for each sample using the Viridot program^4^ The neutralization titers were calculated as follows: 1 – (ratio of the mean number of foci in the presence of sera and foci at the highest dilution of the respective sera sample). Each specimen was tested in duplicate. The FRNT-50 titers were interpolated using a 4-parameter nonlinear regression in GraphPad Prism 9.2.0. Samples that do not neutralize at the limit of detection at 50% are plotted at 20 and used for geometric mean and fold-change calculations. The differences between all groups were determined with the Kruskal–Wallis test with Dunn’s correction for multiple comparisons.

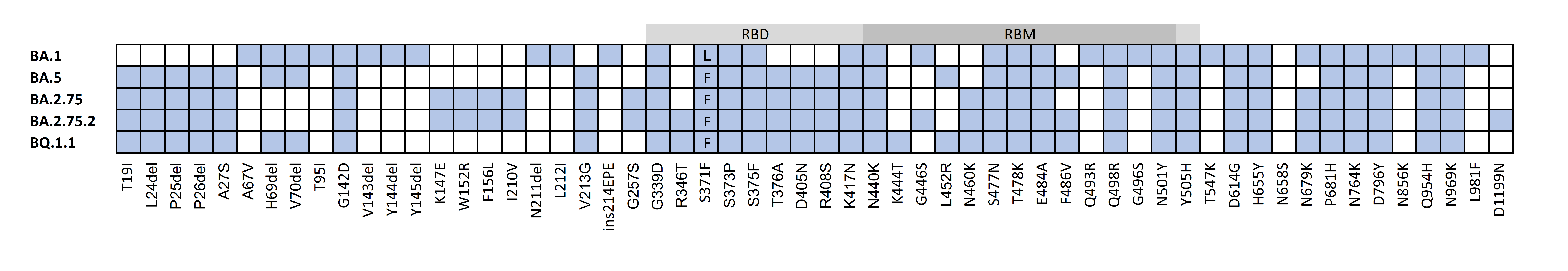

Supplemental Figure 1. Alignment of Spike protein sequence for variants included in this study. White boxes indicate wild-type sequence, colored box indicates amino acid substitution labeled below. Receptor binding domain (RBD) and receptor binding motif (RBM) are indicated above.

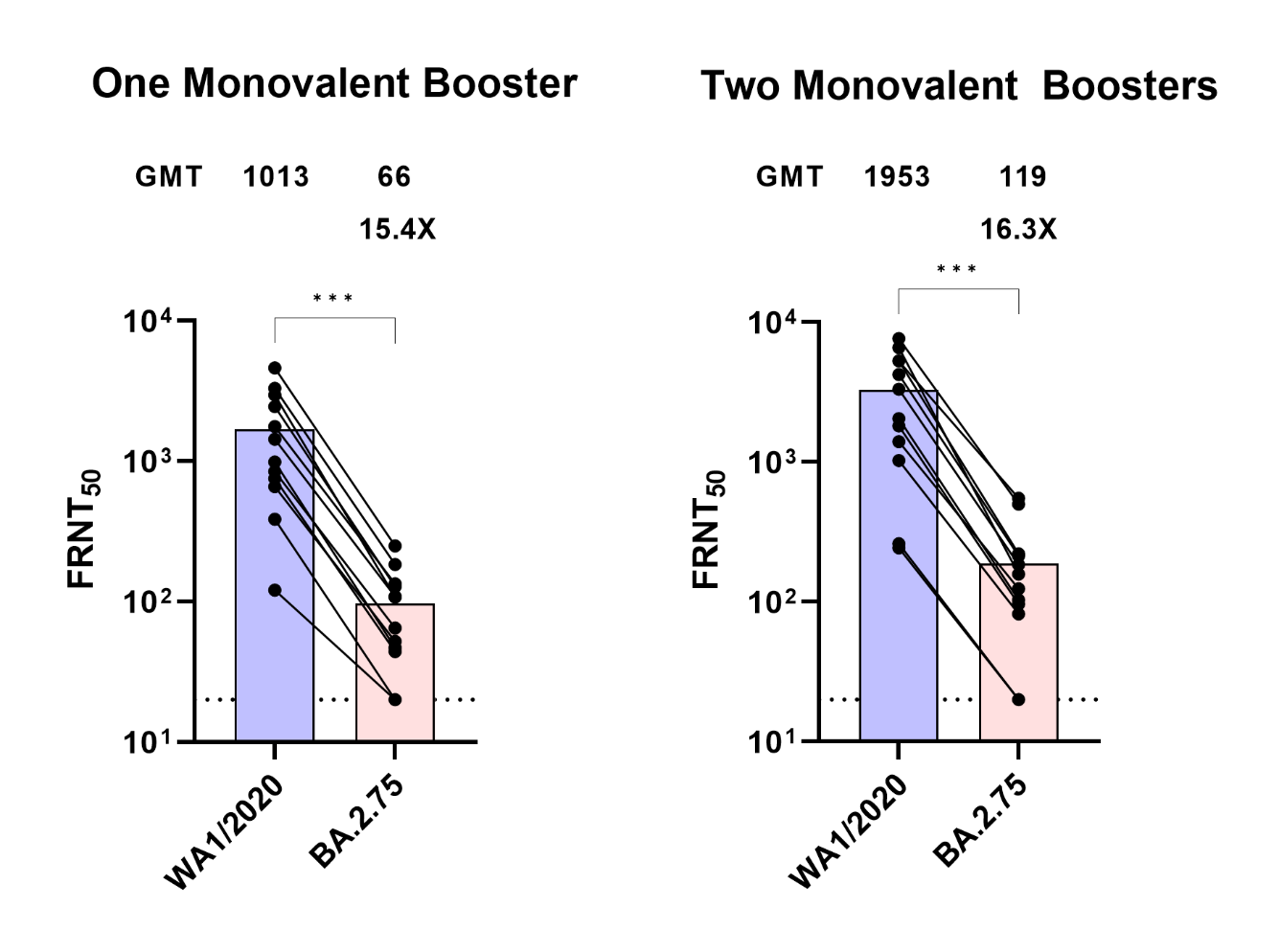

Supplemental Figure 2. Neutralizing responses against WA1, BA.1, BA.5, BA.2.75.2, and BQ.1.1. Shown is the neutralization activity against SARS-CoV-2 variants among 12 individuals who received one monovalent booster (Panel A) and 12 individuals of received two monovalent boosters (Panel B). The focus reduction neutralization test (FRNT_50_ [the reciprocal dilution of serum that neutralizes 50% of the input virus]) geometric mean titers for each variant are shown above each panel along with ratios of GMT compared to WA1/2020. The connecting lines between the variants represent matched serum samples. The horizontal lines represent the limit of detection of the assay (FRNT_50_ GMT 20). *** p<0.0005, Wilcoxon matched-pairs ranked test.

Supplemental Table 1: Amino acid substitutions in spike protein of variants used in the study.

| **Variant** | **Virus Name** | **GISAID** | **Amino Acid Substitutions** |
| --- | --- | --- | --- |
| **BA.1** | hCoV-19/USA/GA-EHC-2811C/2021 | EPI_ISL_7171744 | A67V, H69del, V70del, T95I, G142D, V143del, Y144del, Y145del, N211del, L212I, ins214EPE, S371L, S373P, S375F, G339D, K417N, N440K, G446S, Q493R, Q498R, S477N, T478K, E484A, G496S, N501Y, T547K, Y505H, D614G, H655Y, N679K, P681H, N764K, D796Y, N856K, Q954H, N969K, L981F |
| **BA.5** | hcov-19/USA/MD/HP30386/2022 | EPI_ISL_13512579 | T19I, L24del, P25del, P26del, A27S, H69del, V70del, T76I, G142D, V213G, G339D, S371F, S373P, S375F, T376A, D405N, R408S, K417N, N440K, L452R, S477N, T478K, E484A, F486V, Q498R, N501Y, Y505H, D614G, H655Y, N679K, P681H, N764K, D796Y, Q954H, N969K |
| **BA.2.75** | hCoV-19/USA/CA-Stanford-94_S13/2022 | EPI_ISL_14393635 | T19I, L24del, P25del, P26del, A27S, G142D, K147E, W152R F157L, I210V, V213G, G257S, G339H, S371F, S373P, S375F, T376A, D405N R408S, K417N, N440K, G446S, N460K, S477N, T478K, E484A, Q498R, , N501Y, Y505H, D614G, H655Y, N679K, P681H, N764K, D796Y, Q954H, N969K |
| **BA.2.75.2** | hCoV-19/USA/CA-Stanford-105_S27/2022 | EPI_ISL_15181486 | T19I, L24del, P25del, P26del, A27S, G142D, K147E, W152R, F157L, I210V, V213G, G257S, G339H, R346T, S371F, S373P, S375F, T376A, D405N, R408S, K417N, N440K, G446S, N460K, S477N, T478K, E484A, F486S, Q498R, N501Y, Y505H, D614G, H655Y, N679K, P681H, N764K, D796Y, Q954H, N969K, D1199N |
| **BQ.1.1** | hCoV-19/USA/CA-Stanford-106_S04/2022 | EPI_ISL_15196219 | T19I, L24del, P25del, P26del, A27S, H69del, V70del, G142D, V213G, G339D, R346T, S371F, S373P, S375F, T376A, D405N, R408S, K417N, N440K, K444T, L452R, N460K, S477N, T478K, E484A, F486V, Q498R, N501Y, Y505H, D614G, H655Y, P681H, N679K, N764K, D796Y, Q954H, N969K |

Supplemental Table 2: Demographic information for individuals in single monovalent booster group.

| Sample # | Age | Sex | Exposure status | Vaccine used | Time from vaccination |
| --- | --- | --- | --- | --- | --- |
| 1 | 40 | Male | Naive | Pfizer | 1 week after booster dose |
| 2 | 28 | Male | Naive | Pfizer | 4 weeks after booster dose |
| 3 | 33 | Female | Naive | Pfizer | 4 weeks after booster dose |
| 4 | 31 | Female | Naive | Pfizer | 4 weeks after booster dose |
| 5 | 29 | Female | Naive | Pfizer | 4 weeks after booster dose |
| 6 | 54 | Male | Naive | Pfizer | 4 weeks after booster dose |
| 7 | 51 | Male | Naive | Moderna | 4 weeks after booster dose |
| 8 | 26 | Female | Naive | Moderna | 4 weeks after booster dose |
| 9 | 27 | Male | Naive | Pfizer | 1 week after booster dose |
| 10 | 26 | Female | Naive | Moderna | 4 weeks after booster dose |
| 11 | 33 | Female | Naive | Moderna | 1 week after booster dose |
| 12 | 27 | Female | Naive | Pfizer | 1 week after booster dose |

Supplemental Table 3: Demographic information for double monovalent booster group.

| Sample # | Age | Sex | Exposure status | Vaccine used | Time from vaccination |
| --- | --- | --- | --- | --- | --- |
| 13 | 68 | Female | Naive | Moderna | 100 days after 2^nd^ booster |
| 14 | 72 | Female | Naïve | Moderna | 92 days after 2^nd^ booster |
| 15 | 69 | Male | Naive | Moderna | 92 days after 2^nd^ booster |
| 16 | 72 | Female | Naïve | Moderna | 73 days after 2^nd^ booster |
| 17 | 60 | Female | Naïve | Moderna | 100 days after 2^nd^ booster |
| 18 | 64 | Male | Naïve | Moderna | 83 days after 2^nd^ booster |
| 19 | 71 | Male | Naïve | Moderna | 70 days after 2^nd^ booster |
| 20 | 60 | Male | Naïve | Moderna | 71 days after 2^nd^ booster |
| 21 | 67 | Female | Naïve | Moderna | 71 days after 2^nd^ booster |
| 22 | 62 | Male | Naïve | Pfizer | 54 days after 2^nd^ booster |
| 23 | 53 | Female | Naïve | NA | NA |
| 24 | 70 | Female | Naïve | Moderna | 104 days after 2^nd^ booster |

NA: Data not available

Supplemental Table 4: Demographic information for bivalent booster group.

| Sample # | Age | Sex | Exposure status | Vaccine used | Time from vaccination |
| --- | --- | --- | --- | --- | --- |
| 25* | 70 | F | Naïve | Moderna | 20 days after booster |
| 26 | 47 | M | Naïve | NA | 20 days after booster |
| 27 | 46 | F | Naïve | Moderna | 17 days after booster |
| 28 | 41 | F | Naïve | NA | NA |
| 29 | 36 | F | Naïve | Pfizer | 17 days after booster |
| 30 | 34 | F | Naïve | Moderna | 16 days after booster |
| 31 | 46 | F | Recovered | NA | 28 days after booster |
| 32 | 38 | M | Naïve | NA | 19 days after booster |
| 33** | 23 | M | Naïve | NA | 16 days after booster |
| 34 | 32 | M | Naïve | NA | 17 days after booster |
| 35 | 33 | F | Naïve | NA | 42 days after booster |
| 36 | 40 | M | Recovered | NA | 36 days after booster |

NA: Data not available

*This individual received two monovalent boosters prior to bivalent booster.

**This individual received a primary vaccination of Johnson and Johnson followed by one monovalent booster and one bivalent booster.
